# Ovariectomy and Chemical Ovarian Failure Exacerbate Atherosclerosis without Impairing Limb Recovery in Experimental Peripheral Artery Disease

**DOI:** 10.64898/2026.05.01.722348

**Authors:** Mia Y. Buck, Caroline G. Pass, Jaewon Choi, Divyansha Moparthy, Nathan H. Law, Terence E. Ryan

**Affiliations:** Department of Applied Physiology and Kinesiology; Center for Exercise Science; Myology Institute, The University of Florida, Gainesville, FL, USA

**Keywords:** Atherosclerosis, hindlimb ischemia, ovariectomy, 4-vinylcyclohexene dioxide

## Abstract

**Background:** Peripheral artery disease is a major manifestation of atherosclerotic cardiovascular disease (ASCVD) that affects both men and women. In women, menopause increases the ASCVD risk. However, preclinical ASCVD research has historically been conducted predominantly in males, with relatively few studies focused on females and even fewer incorporating menopause models that more closely reflect human ASCVD pathobiology. Herein, we tested whether the chemical 4-vinylcyclohexene diepoxide (4-VCD)-induced ovarian failure or ovariectomy (OVX) would drive atherosclerotic development and worsen ischemic limb pathophysiology.

**Methods:** Female C57BL/6J mice were injected with adeno-associated virus–mediated encoding a gain-of-function mutant PCSK9 and fed an atherogenic diet for 23 weeks. Based on the baseline body weight, mice were randomly assigned to normally cycling controls (CON), 4-VCD, or OVX groups. Three weeks after the conformation of ovarian failure (4-VCD) or surgical ovarian removal (OVX), hindlimb ischemia (HLI) was induced via femoral artery ligation, and limb perfusion recovery and limb muscle performance were assessed.

**Results:** Both 4-VCD treatment and OVX reduced uterus mass, without impacting body weight or composition, or circulating cholesterol levels compared to CON mice. Despite the similar metabolic and cholesterol profiles, atherosclerotic lesion areas were 1.5–1.7-fold greater in 4-VCD and OVX mice than CON mice. Perfusion recovery following HLI and plantar flexor muscle function in the ischemic limb were similar across groups, though muscle oxygenation was reduced in 4-VCD and OVX groups.

**Conclusions:** Ovarian failure and removal exacerbated atherosclerotic development but had minimal impacts on perfusion recovery and limb function following HLI. These findings confirm the inclusion of menopausal models, whether through ovarian failure or OVX, should be carefully considered to improve translatability of preclinical ASCVD studies, especially for women’s health.

**Clinical Perspective:** *What is New?:* We demonstrate that both gradual ovarian failure (4-VCD) and surgical ovariectomy exacerbate atherosclerotic plaque development in a clinically relevant AAV-PCSK9 model, despite similar circulating lipid levels. In contrast, loss of ovarian function did not impair limb perfusion recovery or muscle functional outcomes following hindlimb ischemia, revealing a dissociation between atherosclerotic burden and limb functional recovery in experimental peripheral artery disease (PAD).

*What are the Clinical Implications?:* These findings provide new insight into why menopause increases atherosclerotic cardiovascular disease (ASCVD) risk while not necessarily demonstrating proportional impairments in limb recovery following ischemia. The data suggest that menopause-associated factors accelerate large-vessel atherosclerosis independent of circulating lipids, highlighting the need for targeted therapies beyond lipid lowering in postmenopausal women. Moreover, the dissociation between plaque burden and ischemic limb function underscores the importance of assessing functional outcomes in PAD independently of vascular imaging. Finally, these findings suggest that the incorporation of menopause-relevant models in preclinical research should be considered within the context of the specific biological endpoints and translational goals being evaluated.

## INTRODUCTION

Peripheral artery disease (PAD) is the third leading cause of atherosclerotic cardiovascular morbidity, making it a significant global health problem^1^. Nearly 236 million people worldwide^2^ and 8 million people in the United States alone^3^ live with PAD. The prevalence of PAD is projected to reach 360 million by 2050 worldwide, affecting 43% and 63% of men and women above 60 years old^4^. Women often present with symptomatic PAD 10 – 20 years later than men^5^, and in many cases with a more severe, advanced disease. Epidemiological studies have reported that less than 2% of women below the age of 50 years old suffer from PAD^6^, but this number dramatically rises to ∼21% and ∼40% in women over the age of 75 and 85 years of age, respectively^5^. Commonly observed between 45 and 55 years of age, menopause is characterized by a decline in estrogen levels and other hormonal shifts. Menopause causes a detrimental increase in systemic low-density lipoprotein (LDL) cholesterol levels that accelerates the development of atherosclerosis^7–9^. As such, postmenopausal women are at higher risk for atherosclerotic cardiovascular disease (ASCVD), including PAD. Despite the growing prevalence of PAD and sex-based disparities in disease progression, the disease remains understudied relative to other forms of ASCVD, particularly in women.

Unfortunately, the vast majority of ASCVD research have focused predominantly on males and has inadequately addressed sex-specific differences^10^. This is particularly evident in preclinical studies, where it is estimated that only 16 – 28% of published atherosclerosis studies have included both male and female animals between 2006 and 2018^11,12^. To raise efforts on integrating knowledge of biological factors in human and vertebrate animal studies, various funding agencies including the American Heart Association^6^ and the National Institutes of Health have issued policies that aimed to address sex as biological variable. Several studies using experimental ischemia models have reported hormone-driven sex differences. For instance, androgen-treated male mice demonstrated greater capillary density and limb perfusion recovery following hindlimb ischemia (HLI)^13^. In females, loss of estrogen exacerbated ischemia-reperfusion injury^14^, while treatment of estrogen increased capillary density^15^, indicating sex-dependent mechanisms of vascular remodeling. Notably, female mice display impaired arteriogenesis and thereby poorer blood flow recovery following HLI compared to males^16,17^. Despite these studies, there is still a substantial gap in knowledge surrounding how biological sex impacts the development of ASCVD and how it may impact the response to subsequent ischemic events. Thus, there is a significant need for the inclusion of both sexes in preclinical studies, yet the optimal conditions for best modeling the human condition remain unclear.

Because ASCVD risk increases dramatically following menopause in women, researchers have surgically removed the ovaries (ovariectomy, OVX) as a way to induce menopause in rodents. However, due to its abrupt hormone loss, this approach does not reflect the menopause transition in humans which involves a gradual decline in ovarian function with residual ovarian tissues retained after full transition. Alternatively, treatment with the chemical 4-vinylcyclohexene diepoxide (4-VCD) has emerged as a menopause model that induces more gradual ovarian failure^18–20^. Daily injection of 4- VCD causes progressive depletion of ovarian follicles that results in a corresponding decrease in estrogen and increase in follicle-stimulating hormone in ovary-intact animals. This gradual transition more adequately reflects the natural progression of human menopause and has been demonstrated to provide a more physiologically relevant means of studying the impacts of sex hormones on the progression of menopause-related diseases^21–23^. While there is evidence showing that both OVX and gradual ovarian failure enhance atherosclerotic burden^24–26^, how these models impact the hindlimb ischemic pathology is unknown.

To address this gap, we aimed to compare the ischemic limb pathology in normally cycling female mice to those that received either OVX or 4-VCD treatment. To better mimic the clinical condition of women with PAD, all mice received a systemic injection of an adeno-associated virus (AAV8) encoding a gain-of-function murine proprotein convertase subtilisin/kexin type9 (PCSK9) mutant and consumed an atherogenic diet throughout the study. We hypothesized that OVX and 4-VCD would worsen the outcomes of limb ischemia and impair recovery following the HLI surgery.

## METHODS

### Animals

All animal procedures were approved by the Institutional Animal Care and Use Committee at the University of Florida (protocol 202300000780). Female C57BL/6J mice were purchased from Jackson Laboratories (stock no. 000664). At 11 weeks old, mice were given a single tail vein injection of murine PCSK9 harboring a D377Y mutation driven by the hcrApoE/hAAT promoter and packaged in the AAV8 serotype to drive expression in the liver (1x10^11^ vg). AAV8-PCSK9 was obtained from Vector Biolabs (Malvern, PA). Immediately following the AAV injection, all mice were placed on an atherogenic diet (45% fat and 0.2% total cholesterol; Envigo Teklad Global, Cat. No. TD.10885) for 23 weeks. Mice were housed in a temperature (22°C) controlled room with 12-h light/dark cycles and *ad libitum* access to food and water. Based on baseline body weight, mice were randomized to control (CON: n=12) that displays normal estrus cyclicity, OVX (n=11), or 4-VCD treatment (n=11).

### 4-VCD Treatment

At 12 weeks of age, mice in the 4-VCD group were given daily intraperitoneal injections of 4-VCD (Millipore-Sigma, Cat. No. 94956) at 160 mg/kg/day prepared in sesame oil (Millipore-Sigma, Cat. No. S3547) for 20 consecutive days. Daily vaginal cytology was performed beginning at 60 days after the first injection to determine cessation of cyclicity as previously described^27^. Mice were considered acyclic after 15 consecutive days without estrous.

### Ovariectomy

A bi-lateral ovariectomy surgery was performed in mice anesthetized via inhaled isoflurane. Pre-operatively, all mice received treatment with slow-release buprenorphine (Ethiqa, 3.25 mg/kg) for analgesia. In a prone position, dorsal midline incision through the skin was made followed by a ∼1-2mm incision through the posterior muscle wall. The oviduct and distal uterine horn were clamped with a hemostat and the ovary was removed. The muscle layer was closed with absorbable sutures and the skin closed with non-absorbable sutures. Post-operatively, mice received subcutaneous injections of meloxicam (15mg/kg) every 24 hours for three days.

### Hindlimb Ischemia

Unilateral femoral artery ligation was performed to induce HLI as previously described^28,29^. Briefly, mice were anesthetized with an intraperitoneal injection of ketamine (90 mg/kg) and xylazine (10 mg/kg), followed by ligation of the femoral artery inferior to the inguinal ligament and immediately proximal to the saphenous and popliteal branches. Mice were given a subcutaneous injection of extended-release buprenorphine (Ethiqa, 3.25 mg/kg) for analgesia.

### Limb Perfusion Imaging

Perfusion recovery following HLI was measured using laser Doppler perfusion imaging (Model LDI2-HIR, Moor Instruments). Mice anesthetized with an intraperitoneal injection of xylazine (10 mg/kg) and ketamine (100 mg/kg) were placed under the scanner in a prone position on a temperature-controlled heating pad. Data were analyzed using MoorLDI Review Software (v6.2)^30^ and were reported as a percentage of the non-ischemic limb.

### Body Composition

Fat and lean body mass, as well as bone density were measured in anesthetized mice using dual-energy X-ray absorptiometry (DEXA InAlyzer 2, Micro Photonics Inc.).

### Blood Measurements

Peripheral blood was collected via tail snip in a heparin-coated capillary tube every four weeks until euthanasia and centrifuged at 1,200g at 4°C for 10 min. Plasma was stored at -80°C until analysis. Blood glucose was measured using OneTouch Ultra test strips and a standard glucometer. Circulating total cholesterol (Cayman Chemical, Cat. No. 10007640) and LDL- and high-density lipoprotein (HDL)-cholesterol (Crystal Chem, Cat. No. 79980 and 79990) levels were measured using a commercially available kit according to the manufacture’s protocol.

### *In-situ* Muscle Function

Muscle function of the plantar flexor complex consisting of gastrocnemius, soleus, and plantaris muscles was assessed *in-situ* using a whole animal system (Aurora Scientific Inc, Model 1300A). Mice were anesthetized with an intraperitoneal injection of xylazine (10 mg/kg) and ketamine (100 mg/kg), and subsequent doses of ketamine (100mg/kg) were given as needed for maintenance. The plantar flexor complex from the ischemic and non-ischemic limbs was isolated from its distal insertion while leaving all vasculature intact. The distal portion of *Achilles* tendon was tied with a 4-0 silk suture attached to the lever arm of the force-length servomotor. The sciatic nerve was isolated and stimulated at 2mA via bipolar electrodes using square-wave pulses (Aurora Scientific, model 701A stimulator) to elicit muscle contractions. Lab-View–based DMC program (version 615A.v6.0, Aurora Scientific Inc) was used for data collection and servomotor control. Muscle function testing was performed on a temperature-controlled platform to maintain a body temperature of 37°C.

First, optimal length was obtained via twitch contractions. Next, isometric contractions were elicited at stimulation frequencies of 1Hz, 40Hz, 80Hz, and150Hz (2 mA; 0.2-ms pulse width; 500-ms train duration) with one minute of rest between contractions. Following a three-minute recovery period, we performed a 6-minute limb function test developed for preclinical models of HLI^31^ to assess muscle performance, perfusion flux, and oxygenation in response to a sequence of after-loaded isotonic contractions. Briefly, the plantar flexor complex was stimulated at 80Hz while the lever arm was allowed to shorten when force exceeded 30% of the absolute force at the 80Hz isometric contraction. This isotonic (shortening) contraction was followed by a 2.5-second static rest period to allow perfusion and oxygenation measurements in the gastrocnemius muscle and then three small passive stretches. This protocol was repeated every ∼4 seconds for six minutes. Data from the 6-minute limb function test were analyzed using custom scripts in MATLAB (MathWorks). The average perfusion flux and total hemoglobin, oxyhemoglobin, and deoxyhemoglobin levels for each motionless rest period were quantified and plotted over time. Power, shortening velocity, and mechanical work were calculated for every contraction and plotted across time. The total work performed was calculated by summing the work of each individual contraction.

### Labeling of Perfused Capillaries

Mice were given a retro-orbital injection of 50 µL *Griffonia simplicifolia* lection isolectin B4, Dylight 649 (Vector Laboratories, Cat. No. DL-1208) 1mg/mL to fluorescently label α-galactose residues on the surface of endothelial cells of perfused capillaries. Animals were returned to their cage and allowed 1-2 hours of free movement to permit systemic circulation of the labeled isolectin prior to the 6-minute limb function test and subsequent tissue harvesting.

### Immunofluorescence Microscopy

The medial gastrocnemius muscle was carefully dissected, embedded on a cryomold with optimal cutting temperature compound, and frozen in liquid nitrogen-cooled isopentane. Transverse sections were cut from the muscle at 10µm using a Leica 3050S cryostat and mounted on microscope slides. All muscle sections were then blocked at room temperature for one hour in 1x phosphate-buffered saline (PBS) containing 5% goat serum and 1% bovine serum albumin. To quantify total capillaries, sections were stained with an anti-CD31 primary antibody (1:100, Abcam, Cat. No. ab28364) and Alexa Fluor 488 goat anti-rabbit immunoglobulin G (1:250, ThermoFisher Scientific, Cat. No. A11008). Arterioles were labeled with alpha smooth muscle actin (1:400, ThermoFisher Scientific, Cat. No. 14976082) and Alexa Fluor 555 goat anti-mouse immunoglobulin G2a (1:250, ThermoFisher Scientific, Cat. No. A21137), along with rabbit anti-laminin (1:200, Millipore-Sigma, Cat. No. L9393) and Alexa Fluor 488 goat anti-rabbit immunoglobulin G to visualize the basal lamina surrounding myofibers. Mouse anti-CD68 (1:100, ThermoFisher Scientific, Cat. No. 14-0688-82) and Alexa Fluor 488 goat anti-mouse immunoglobulin G1 (1:250, ThermoFisher Scientific, Cat. No. A21121) were used to stain macrophages. On the same slide, M1 macrophages were visualized with Armenian hamster anti-CD80 (1:200, ThermoFisher Scientific, Cat. No. 14-0801-82) and Alexa Fluor 555 goat anti-Armenian hamster immunoglobulin G (1:250, ThermoFisher Scientific, Cat. No. A78964). All primary antibodies were applied overnight at 4°C. The following day, slides were washed in 1× PBS and all secondary antibodies or directly conjugated dyes were applied for 1.5 hours at room temperature. Coverslips were mounted using Vectashield hardmount with DAPI (Vector Laboratories, Cat. No. H-1500). All slides were imaged using an Evos FL2 Auto microscope (ThermoFisher Scientific) at 20x magnification. Obtained images were thresholded, and the number of perfused capillaries, total capillaries, arterioles, pan macrophages, and M1 macrophages were quantified using a particle counter in Fiji/ImageJ (National Institutes of Health). Skeletal myofiber cross-sectional area (CSA) was quantified using MuscleJ2, an automated analysis software developed in Fiji^32^.

### Quantification of Atherosclerotic Lesions

The thoracic aorta was carefully dissected as previously described^33^ and rinsed with 1x PBS for one minute in a 35mm petri dish. Tissue was then submerged in 85% propylene glycol for two minutes and incubated in filtered Oil-Red-O (StatLab, Cat. No. STORO100) for six minutes, followed by a one-minute wash with 85% propylene glycol, two five-minute washes with 0.1% PBST, and a one-minute wash with 1x PBS. Aortas were then transferred to a microscope slide and imaged using MU Series 10.0MP USB 2.0 Color CMOS C-Mount Microscope Camera (AmScope) at 17.5x magnification. Atherosclerotic lesion was quantified as the proportion of Oil-Red-O positive area relative to the total aorta area in ImageJ.

### Statistical Analysis

All data are presented as mean ± standard deviation. Normality was determined by the Shapiro-Wilk test. Data with normal distribution were analyzed using one-way or a two-way analysis of variance with Tukey’s post hoc multiple comparisons where appropriate. Statistical analyses were performed using GraphPad Prism software (version 10.5.0). Statistical significance was declared at *P* < 0.05.

## RESULTS

### Treatment of 4-VCD and OVX Reduced Uterus Weight with No Impact on Body Weight and Composition

To test whether chemical-induced ovarian failure or OVX exacerbates atherosclerotic development and ischemic pathophysiology, mice were given a single injection of AAV8-PCSK9 and fed an atherogenic diet (**Figure 1A**). Menopause was confirmed at 118 ± 12 days following the onset of 4-VCD injection. To match closely the duration of ovarian failure, we performed OVX surgeries at week 16 of the intervention, which equates to ∼112 days after the AAV8-PCSK9 and atherogenic diet treatment. As expected, 4-VCD and OVX mice had significantly smaller uterus masses compared to normally cycling females (**Figure 1B**), confirming an estrogen depletion. Unsurprisingly, consumption of the atherogenic diet resulted in a progressive increase in body weight in all groups without any statistical differences across groups (**Figure 1C**). We observed an anticipated decrease in body weight following HLI surgery but was not affected by the menopause status. All groups demonstrated a significant increase in the percentage of body fat across time (*P*=1.170E-47). Interestingly, 4-VCD mice demonstrated an early increase in body fat by four weeks and also displayed the most pronounced (2.4-fold) increase at 16 weeks compared to baseline (*P*=0.0001), with no impact of HLI. As expected, the progressive increase in body fat mass was accompanied by a progress decrease in the percentage of lean mass (*P*=3.626E-50), which was observed in all groups (**Figure 1C**). Despite being randomized by baseline body weights, mice assigned to the OVX group displayed consistently lower percent fat mass and higher percent lean masses, although these were not different from other groups with posthoc testing (*P*>0.09 for all time points). Bone mineral density was lower in OVX mice than CON mice both before and after HLI. This suggests that OVX, but not 4-VCD induced menopause, had a significant impact on bone health in this experimental paradigm.

**Figure 1.**
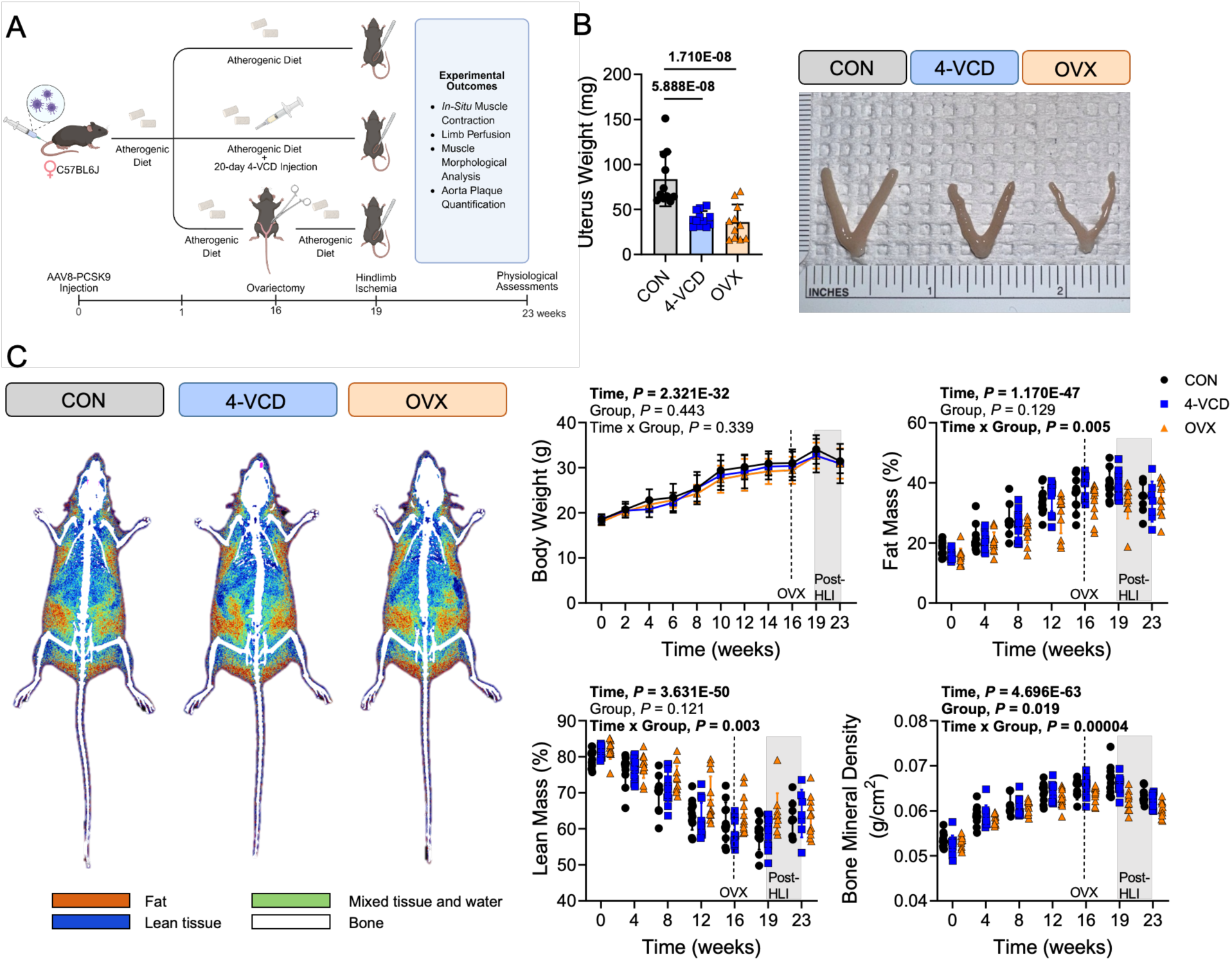
Chemical-induced ovarian failure and ovariectomy decreases uterus size but do not impact body composition. (A) Graphic of the experimental design and study timeline (generated using BioRender). (B) Mass and representative images of uterus. Uterus weight was analyzed using one-way ANOVA. (C) Body mass, composition, and bone mineral density were measured via dual-energy x-ray absoptiometry and analyzed using two-way ANOVA. Error bars represent the standard deviation. All panels contain n=11 (CON), n=12 (4-VCD), and n=11 (OVX).

### Impacts of Ovarian Failure on Blood Glucose, Cholesterol, and Atherosclerosis

Consumption of the atherogenic diet significantly increased blood glucose concentrations in all groups (**Figure 2A**), although this reached a steady state at four weeks, and no group differences were observed (*P*=0.24). All mice displayed a drastic increase in total cholesterol levels (**Figure 2B**); however, HLI resulted in a ∼50% reduction in total cholesterol that may be related to decreased food intake. Plasma LDL cholesterol levels, a primary, casual factor for atherosclerosis, increased by 5.4-fold at eight weeks into the diet intervention (68.83±1.25 vs. 367.33±13.96 mg/dL) and plateaued for the duration of the study (**Figure 2C**). Similarly, the atherogenic diet elevated HDL cholesterol levels in all groups but was not impacted by menopause status (**Figure 2D**). In contrast, atherosclerotic plaque lesion area in the thoracic aorta was 1.7–fold and 1.5 –fold greater in 4-VCD and OVX groups compared to the normally cycling CON mice (**Figure 2E**). These findings indicate that both 4-VCD and OVX exacerbate atherosclerosis despite having similar circulating cholesterol levels while consuming the atherogenic diet.

**Figure 2.**
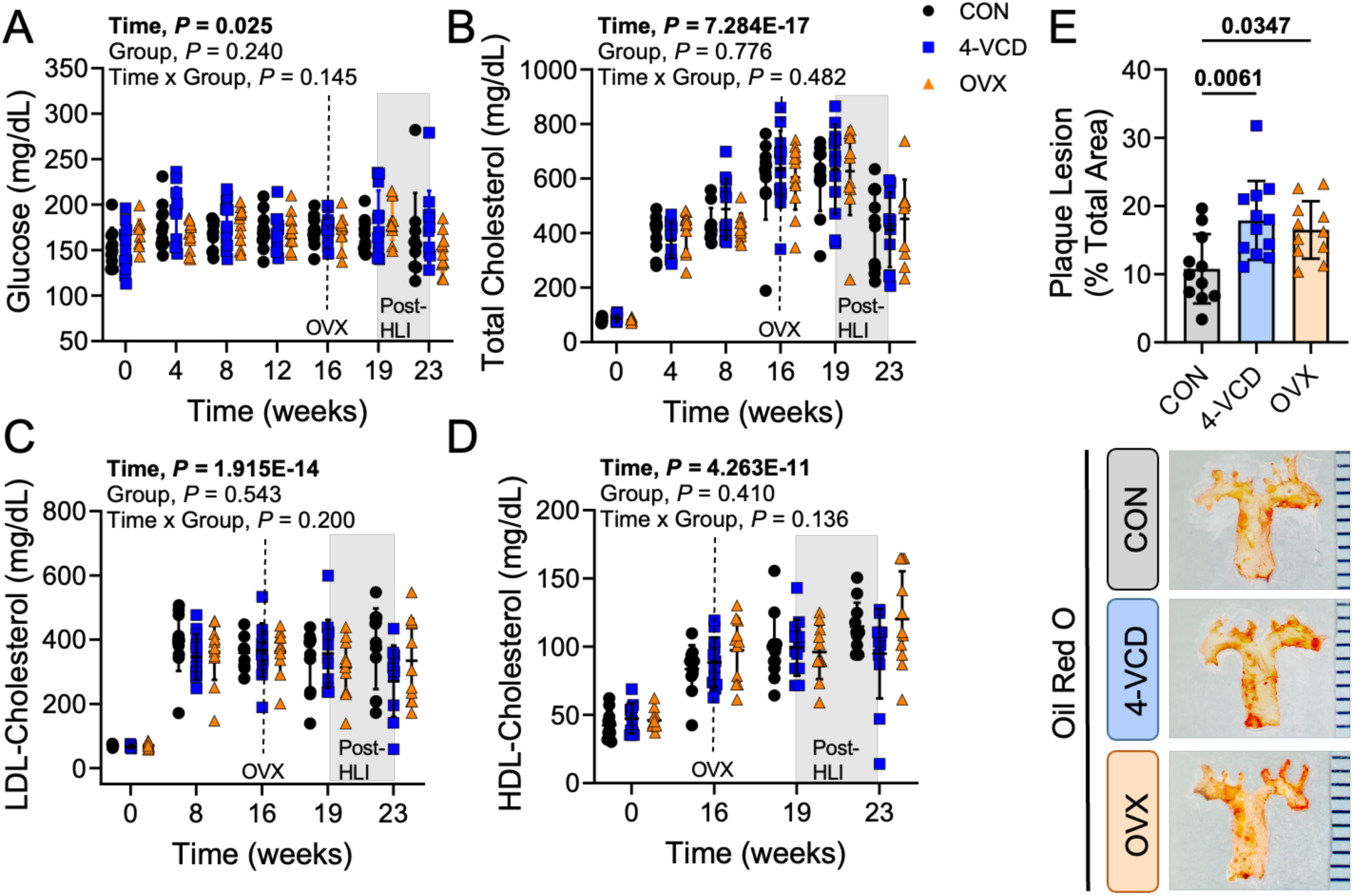
Impacts of Ovarian Failure on Blood Glucose, Cholesterol, and Atherosclerosis. (A) Quantification of whole blood glucose and plasma total, LDL-, and HDL- cholesterol levels throughout the study. Data were analyzed using two-way ANOVA. (B) Quantification of atherosclerotic lesion, expressed as percentage of total aorta area, and representative microscopic images. Data were analyzed using one-way ANOVA. Error bars represent the standard deviation. All panels contain n=11 (CON), n=12 (4-VCD), and n=11 (OVX).

### Neither 4-VCD nor OVX Altered Limb Perfusion Recovery Following HLI

To test whether 4-VCD and OVX models have an impact on perfusion recovery, laser Doppler perfusion imaging was performed on all mice before and after HLI surgery. While perfusion recovered gradually over time (*P*=2.926E-16), no group effects were observed (**Figure 3A**), indicating that ovarian failure or removal did not negatively impact perfusion recovery in mice. Correspondingly, total and perfused capillaries, as well as the number and size of arterioles were similar across groups (**Figure 3B**).

**Figure 3.**
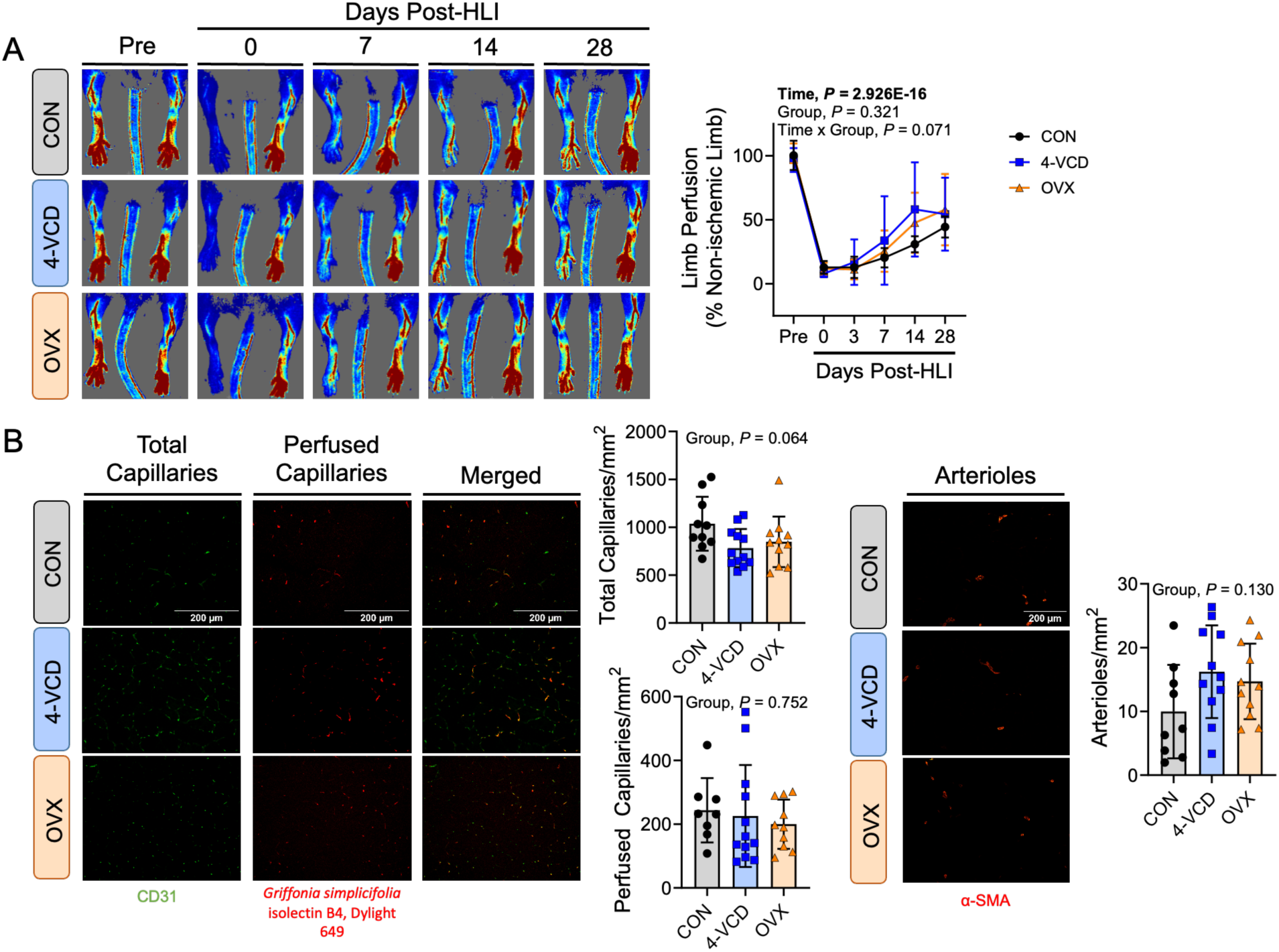
Neither 4-VCD nor OVX Altered Limb Perfusion Recovery Following HLI. (A) Laser Doppler perfusing imaging quantification of perfusion recovery expressed as percentage of control limb. Data were analyzed using two-way ANOVA (B). Quantification and representative images of total and perfused capillaries and arterioles. Data were analyzed using one-way ANOVA. Error bars represent the standard deviation. Panel A contains n=11 (CON), n=12 (4-VCD), and n=11 (OVX). Panel B contains n=8 (CON), n=11 (4-VCD), and n=10 (OVX).

### Ischemic Limb Function is Not Impaired by 4-VCD or OVX Treatment

Calf muscle pathology is a hallmark of PAD that contributes to the limited walking capacity and mobility impairments seen in humans^34^. To test whether 4-VCD or OVX treatments impaired ischemic muscle function, we performed an assessment of isometric muscle strength and a 6-minute limb function test^31^ in the HLI limb of mice. In the HLI limb, we did not observe any effect of menopause status on plantar flexor muscle mass (**Figure 4A**) or gastrocnemius muscle myofiber area (**Figure 4B**). Plantar flexor muscle strength (absolute force) was also not different between groups (**Figure 4C**). When the absolute force was normalized to muscle weight (specific force), which assesses muscle quality, 4-VCD mice exhibited greater specific force than CON mice (**Figure 4C**). Next, we employed a 6-minute limb function test to quantify power and work outputs in the plantar flexor muscle across a series of repeated contractions. Similar to muscle specific force, 4-VCD and OVX mice exhibited slightly higher muscle power and work outputs across the 6-minute limb function test (**Figure 4D**). However, quantification of the total work performed, akin to the distance covered by PAD patients during a 6-minute walk test, was not different between groups (*P*=0.63, **Figure 4E**). Interestingly, plantar flexor perfusion flux was higher in mice treated with 4-VCD (**Figure 4F**). Notably, plantar flexor oxygenation was significantly lower in both 4-VCD and OVX mice when compared to CON mice (**Figure 4G**). These observations demonstrated that while ovarian failure or removal may have modest effects on active muscle perfusion and oxygenation, work output in the ischemic limb is unaffected. Like the ischemic limb, mice treated with 4-VCD or OVX had poorer oxygenation outcomes in the contralateral non-ischemic limb when compared to normally cycling CON mice, and there were no differences in muscle work output (**Figure 5**).

**Figure 4.**
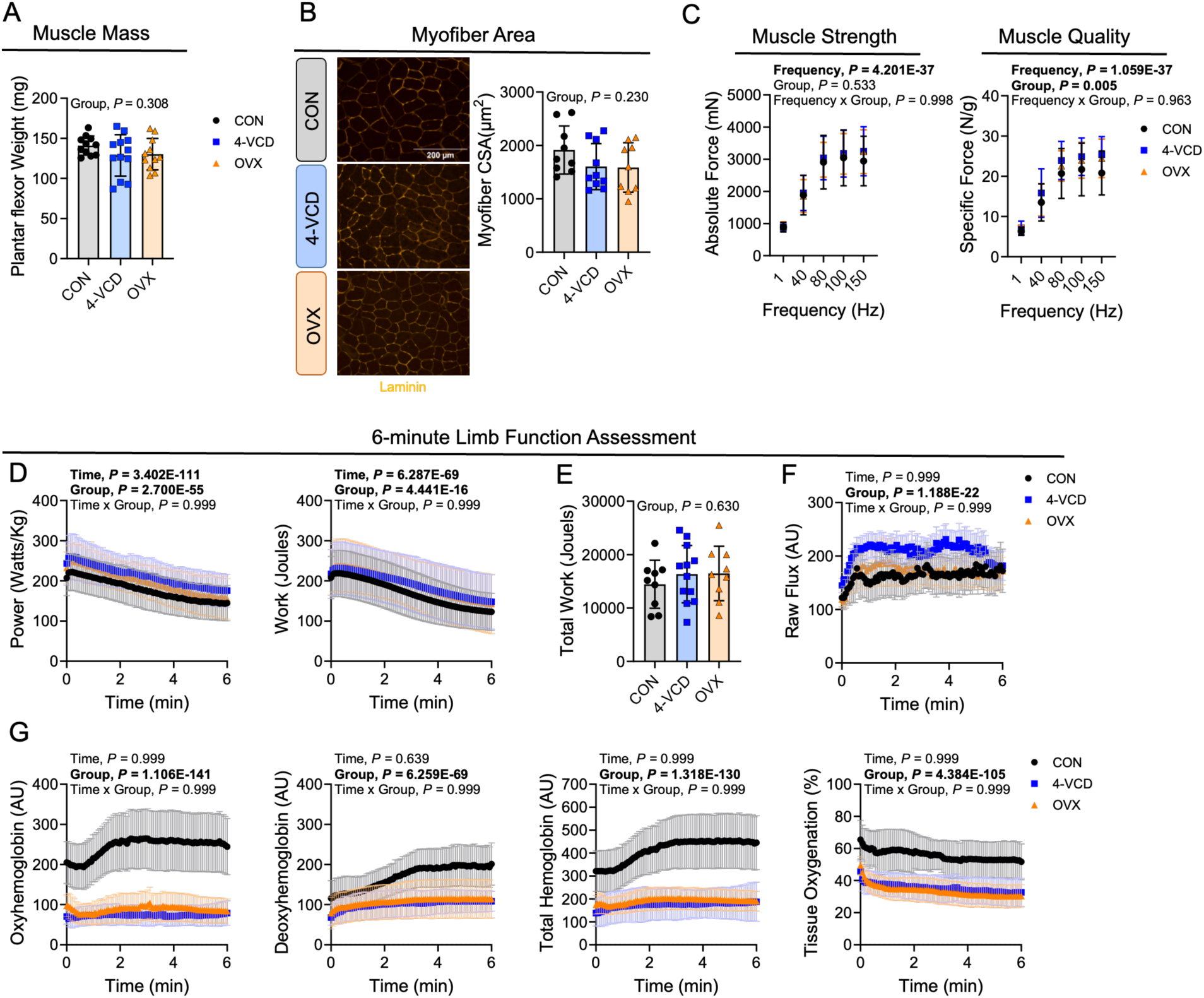
Ischemic Limb Function is Not Impaired by 4-VCD or OVX Treatment. (A) Plantar flexor muscle mass analyzed using one-way ANOVA. (B) Immunofluorescence images and quantification of myofiber cross sectional area (CSA) analyzed using one-way ANOVA. (C) Plantar flexor muscle strength (absolute force) and quality (specific force) analyzed using one-way ANOVA. (D) Muscle power and work output during the 6-minute limb function assessment analyzed using two-way ANOVA. (E) Quantification of total work performed across the 6-minute limb function test analyzed using one-way ANOVA. (F) Perfusion flux measured via laser Doppler flowmetry during the 6-minute limb function test. (G) Muscle oxygenation across the 6-minute limb function test analyzed using two-way ANOVA. Error bars represent the standard deviation. Panel A contains n=11 (CON), n=12 (4-VCD), and n=11 (OVX). Panels B – G contains n=9 (CON), n=12 (4-VCD), and n=9 (OVX).

**Figure 5.**
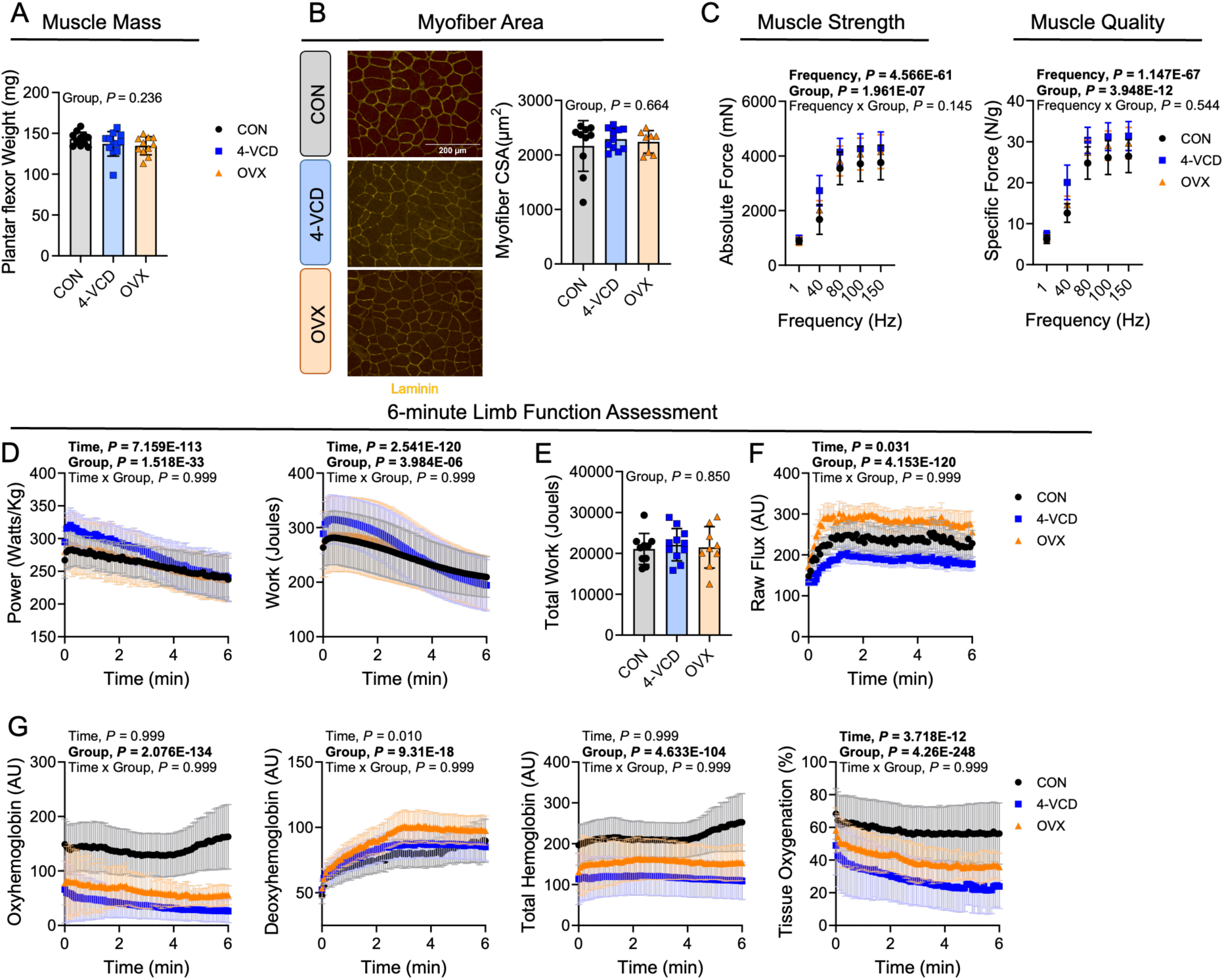
Non-Ischemic Control Limb Function is Not Altered by 4-VCD or OVX Treatment. Plantar flexor muscle mass of the non-ischemic control limb analyzed using one-way ANOVA. (B) Immunofluorescence images and quantification of myofiber cross sectional area (CSA) analyzed using one-way ANOVA. (C) Plantar flexor muscle strength (absolute force) and quality (specific force) analyzed using one-way ANOVA. (D) Muscle power and work output during the 6-minute limb function assessment analyzed using two-way ANOVA. (E) Quantification of total work performed across the 6-minute limb function test analyzed using one-way ANOVA. (F) Perfusion flux measured via laser Doppler flowmetry during the 6-minute limb function test. (G) Muscle oxygenation across the 6-minute limb function test analyzed using two-way ANOVA. Error bars represent the standard deviation. Panel A contains n=11 (CON), n=12 (4-VCD), and n=11 (OVX). Panel B contains n=10 (CON), n=11 (4-VCD), and n=8 (OVX). Panel C – G contain n=10 (CON), n=11 (4-VCD), and n=9 (OVX).

## DISCUSSION

Loss of ovarian hormones after menopause has been widely recognized as an important modifier of ASCVD risk and progression in women. Despite the well-documented sex-dependent differences in ASCVD pathophysiology, females have been underrepresented in both clinical and preclinical research, and there remains no standardized approach for modeling menopause in female rodents. Herein, we employed 4-VCD and OVX models to investigate how ovarian failure and surgical ovarian removal influence atherosclerotic development and limb functional outcomes following an ischemic injury in female mice. The results demonstrated that both 4-VCD and OVX markedly increased the atherosclerotic lesion area compared to normally cycling CON mice despite the similar circulating lipid profiles across groups. To determine whether the loss of ovarian function impacts ischemic limb pathobiology, we next subjected mice to HLI. Rigorous assessments of hindlimb function demonstrated that the loss of ovarian function had no significant impact on ischemic limb work or power output, nor perfusion recovery, though it did result in modestly decreased muscle oxygenation during contraction.

Historically, OVX has been the most commonly used model to study diseases relevant to postmenopausal women, including ASCVD. Ovarian removal causes an abrupt loss of estrogen, resulting in reduced hepatic mRNA expression of LDL receptors and PCSK9^35^ and therefore elevated circulating LDL cholesterol^36,37^. In contrast to OVX, 4-VCD induces menopause through progressive depletion of ovarian follicles while leaving the ovaries intact. Despite the different physiological mechanisms, female animals treated with 4-VCD display impaired lipid metabolism similar to OVX when combined with additional metabolic stressors such as a high-fat diet^38^. In support of this, Mayer *et al.* reported that LDL receptor-deficient female mice treated with 4-VCD or OVX exhibited similar total cholesterol levels and atherosclerotic lesion areas; however, the menopause status did not further exacerbate atherosclerotic burden compared to normally cycling mice^26^. Herein, circulating total, LDL-, and HDL-cholesterol levels were similar between 4-VCD, OVX and normally cycling CON mice, while there was a ∼1.7-fold increase in the atherosclerotic plaque lesion in 4-VCD and OVX mice. This result is consistent with previous studies. For example, Bourassa *et al*. reported that atherosclerotic lesion areas were increased in apolipoprotein E-deficient female mice with OVX compared to normally cycling mice despite similar LDL cholesterol levels^24^. These authors also demonstrated that treatment with 17β-estradiol lowered cholesterol levels and atherosclerotic lesions. Marsh *et al*. reported identical findings in LDL receptor knockout mice with OVX and exogenous estradiol treatment^39^. Two subsequent studies provided evidence that the atheroprotective effects of estradiol are mediated by the estrogen receptor alpha^40,41^. It is important to note that estradiol dosing can impact the degree of atheroprotection as one study reported that while higher doses are protective, lower doses of estradiol can enhance atherosclerosis in ApoE deficient female mice with OVX^42^. Despite the increasing adoption of the AAV-PCSK9 model to study atherosclerosis across diverse biological contexts, the role of ovarian function and menopause has remained largely unexplored in this model. To our knowledge, our study is one of the first to confirm that ovarian failure or removal exacerbates atherosclerosis in the AAV8-PCSK9 atherosclerosis model, confirming previous studies in other atherosclerosis models showing that the plaque burden is influenced by ovarian function even when differences in circulating lipids are absent.

It has been well documented that women face much higher risk of PAD at a later stage in life and often present clinically with more advanced disease than men^43,44^. Unfortunately, only a few preclinical studies have examined the impact of biological sex on ischemic limb pathology, and far less is known about whether ovarian failure impacts further effects. Peng *et al*. reported that healthy female mice displayed poorer perfusion recovery following HLI than males, which was accompanied by reduced numbers of total capillaries and arterioles in the hindlimb muscle^17^. Another study found that androgen treatment rescued reduced blood perfusion recovery following HLI in orchectomzied male mice, suggesting sex hormones do impact ischemic outcomes. In females, estrogen has been shown to promote vasodilation by increasing nitric oxide production from endothelial cells, thereby increasing blood flow^45^. Accordingly, an intramuscular injection of estrogen increased limb blood pressure and capillary to muscle fiber ratio following an ischemic injury in ovariectomized female rabbits^15^. In contrast, estrogen deficiency decreased blood perfusion and capillary density, along with lower protein levels of endothelial nitric oxide synthase in female mice with HLI^46^. In the present study, we found that blood recovery perfusion and total and perfused capillaries in response to HLI were unaffected by chemically-induced or surgical ovarian failure. However, our 6-minute limb function test revealed that 4-VCD and OVX mice demonstrated reduced muscle oxygenation compared to CON mice, though there were no effects on work output. This observation suggests that although 4-VCD and OVX mice were able to preserve their ischemic muscle function, this may have been achieved by increasing muscle deoxygenation alongside slight increases in hyperemia, indicating the presence of altered energetic or mechanical efficiency.

The current study has some limitations. First, although the 4-VCD model offers a more physiologically relevant approach to mimic human menopause compared to OVX, there is a wide variability in the time required for mice to reach ovarian failure. In previous cyclicity studies, ovarian failure had been reported to occur around 54 – 80 days following 20 consecutive injections of 4-VCD at 160 mg/kg/day in healthy female mice maintained on normal chow diet^18,23^. Contrary to this, we observed menopause at 118 ± 12 days following the onset of 20-day injections. This discrepancy is likely due to differences in the experimental design; we combined the 4-VCD treatment with a single injection of PCSK9 and atherogenic diet, which resulted in a significant increase in body weight and circulating cholesterol levels. These metabolic alterations may have delayed or slowed follicle depletion. Accordingly, the length of the dosing should be carefully considered based on a study design and animal model employed to optimize the length of perimenopause. We also acknowledge that the duration of the study following OVX or the confirmation of menopause following 4-VCD treatment may not have captured longer-term effects of estrogen depletion on atherosclerotic development and ischemic limb outcomes. Herein, the substantial increase in LDL cholesterol, a key driver of atherosclerosis, was observed in all groups at eight weeks into the diet intervention, with no further increase for the remaining of the study. In agreement, Ramírez-Hernández *et al*., reported no differences in serum LDL cholesterol levels between 10-week and 20-week post-OVX in rats^47^. This suggests that alternations in lipid metabolism following ovarian failure or removal may plateau over time rather than progressively worsen. Another limitation of this study is the use of relatively young mice and, unfortunately, the availability and cost of aged mice remains a barrier for preclinical researchers. It is plausible that the addition of aging biology may be a significant modifier of the ASCVD and ischemic limb pathobiology studied herein.

In summary, the current study demonstrated both chemically-induce ovarian failure and surgical ovarian removal significantly and equally increase atherosclerotic lesion area. In contrast to our hypothesis, ischemic limb pathology and function were unaffected by ovarian failure. Nonetheless, we suggest that researchers carefully consider the benefits and drawbacks of the inclusion of menopausal female models, whether through chemical-induced ovarian failure or surgical ovary removal, in preclinical studies focused on the pathogenesis of both ASCVD and PAD.

## Non-standard Abbreviations and Acronyms

4-VCD 4-vinylcyclohexene diepoxide

AAV adeno-associated virus

ASCVD atherosclerotic cardiovascular disease

CSA cross sectional area

HDL high-density lipoprotein

HLI hindlimb ischemia

LDL low-density lipoprotein

OVX ovariectomy

PAD peripheral artery disease

## Acknowledgements

None.

## Sources of Funding

This study was supported by the National Institute of Health grants R01-HL171050 and R01-HL149704, as well as American Heart Association grant 25EIA1369187 (TER). CGP supported by the American Heart Association grant 24PRE1196311.

## Disclosures

The authors report no conflicts of interest.

## Notes

### Competing Interest Statement

The authors have declared no competing interest.

